# Time between milestone events in the Alzheimer’s disease amyloid cascade

**DOI:** 10.1101/2020.05.18.103226

**Authors:** Philip S. Insel, Michael C. Donohue, David Berron, Oskar Hansson, Niklas Mattsson-Carlgren

## Abstract

**Objective:** Estimate the time-course of the spread of key pathological markers and the onset of cognitive dysfunction in Alzheimer’s disease.

**Methods:** In a cohort of 336 older adults, ranging in cognitive functioning, we estimated the time of initial changes of Aβ, tau, and decreases in cognition with respect to the time of Aβ-positivity.

**Results:** Small effect sizes of change in CSF Aβ42 and regional Aβ PET were estimated to occur several decades before Aβ-positivity. Increases in CSF tau occurred 11-12 years before Aβ-positivity. Temporoparietal tau PET showed increases 4-5 years before Aβ-positivity. Subtle cognitive dysfunction was observed 7-9 years before Aβ-positivity.

**Conclusions:** Increases in tau and cognitive dysfunction occur years before the presence of significant Aβ. Explicit estimates of the time for these events provide a clearer picture of the time course of the amyloid cascade and identify potential windows for specific treatments.

## Introduction

Disconcerting clinical trial results for the treatment of Alzheimer’s disease (AD) have led to a shift toward earlier intervention, focusing on the early clinical or presymptomatic phases, when biomarkers are needed to identify the disease. The amyloid cascade (Hardy and Selkoe, 2002) is thought to start with elevated levels of two key amyloids in the brain, β-amyloid (Aβ) and tau, and end with severe cognitive and functional impairment (Jack et al., 2010). Growing evidence suggests that an early sign that the cascade has begun is change in cerebrospinal fluid (CSF) Aβ, potentially detectable prior to significant Aβ deposition in the brain as measured by positron emission tomography (PET) (Palmqvist et al., 2016). This accumulation of Aβ has been suggested to be followed by increases in CSF tau and the spread of tau pathology beyond the temporal lobe (Braak and Braak, 1991; Schöll et al., 2016). The build-up and spread of these two brain pathologies is paralleled by gradual cognitive and functional decline (Zetterberg and Mattsson, 2014).

Previous neuropathological and biomarker data suggest that the overall time course of AD is several decades (Li et al., 2017; Villemagne et al., 2013). In autosomal dominant AD, the estimated years to clinical onset has been used to estimate the timecourse of different biomarkers in AD (Bateman et al., 2012). However, the time-course of the spread of Aβ and tau and the onset of clinical symptoms in sporadic AD is unknown. With repeated measures of Aβ over time, the level and rate of change with respect to the key initiating AD pathology may offer a measure of disease progression in sporadic AD. With level and change information, the time from the threshold for significant Aβ pathology can be estimated within individuals, providing the temporal disease progression information important for evaluating biomarker trajectories. Without longitudinal information, cross-sectional studies frequently categorize subjects into two groups – those below a threshold for significant pathology and those above, where subjects just below the threshold who will cross over within months are considered pathologically equivalent to subjects who will not cross over for decades. By incorporating longitudinal information, disease progression with respect to Aβ pathology can be represented to reflect its continuous nature, resulting in a more powerful way to model the relationship between Aβ and downstream processes.

The aim of this study was to evaluate time-from-Aβ-positivity (TFAβ+) in sporadic AD. Using serial 18F-florbetapir (Aβ) PET measurements, rates of change of Aβ were estimated and used to calculate the time-from-threshold for each subject. These subject-specific estimates of the proximity to the threshold for Aβ-positivity (Aβ+) were then used to model the trajectories and temporal ordering of other key markers in AD including CSF Aβ42, regional Aβ PET, several measures of tau including CSF phosphorylated (P-tau) and total tau (T-tau), regional 18F-flortaucipir (AV-1451) tau PET, and cognition. Estimates of the time and ordering of these pathophysiological changes may facilitate the design of future prevention trials and identify a window for early treatment.

## Materials and methods

### Standard protocol approvals, registrations, and patient consents

This study was approved by the Institutional Review Boards of all of the participating institutions. Informed written consent was obtained from all participants at each site.

### Data Availability

All data is publicly available (http://adni.loni.usc.edu/).

### Participants

Data were obtained from the Alzheimer’s Disease Neuroimaging Initiative (ADNI) database (http://adni.loni.usc.edu/, www.adni-info.org) on 1/21/2020. An initial analysis was done on all ADNI participants with available Aβ PET data (in N=963 CU, Aβ+ MCI and Aβ+ AD), to facilitate the estimation of TFAβ+. The population in the primary analysis only included ADNI participants with measurements of both Aβ and tau PET. Of these, all cognitively unimpaired (CU), prodromal AD (Aβ+ MCI) and Aβ+ AD dementia participants were included in the analysis, where Aβ-positivity was defined using a previously established threshold (Standardized Uptake Value Ratio, SUVR = 1.10) (Joshi et al., 2012). Aβ-MCI (N=224, including Aβ-CU to MCI progressors) and Aβ-“AD dementia” subjects (N=51, including Aβ-MCI to AD dementia progressors; we consider these to be misdiagnosed, because we assume AD requires Aβ+) were not included in the main analysis given our aim to model disease progression over the AD continuum and not other diseases, but visualizations of their biomarker data are included for comparison in Figures 2–4 (see Figure legends). Additional description is included in the statistical analysis section.

### Cerebrospinal fluid biomarker concentrations

Cerebrospinal fluid (CSF) samples were collected at baseline by lumbar puncture in a subsample (N=185). CSF Aβ42, total tau (T-tau) and phosphorylated tau (P-tau) were measured by an xMAP assay (INNOBIA AlzBio3, Ghent, Belgium, Fujirebio), as described previously (Olsson et al., 2005; Shaw et al., 2009).

### PET Imaging

Methods to acquire and process Aβ (18F-florbetapir) PET image data were described previously (Landau et al., 2012). We used an a priori defined threshold for Aβ-positivity (SUVR=1.1) (ADNI, 2012; Joshi et al., 2012) applied to the ratio of the average of the four target regions (temporal, cingulate, frontal, and parietal lobes) and the cerebellum, in the estimation of time-from-Aβ-positivity, described in detail below. In a second part of the analysis, five Aβ PET ROI outcomes were considered (Landau and Jagust, 2015; Mormino et al., 2009), (1) the temporal lobe (middle and superior temporal lobe), (2) the parietal lobe (precuneus, supramarginal, inferior and superior parietal lobe), (3) the cingulate gyrus (isthmus, posterior, caudal and rostral anterior cingulate), (4) the frontal lobe (pars opercularis, pars triangularis, pars orbitalis, caudal/rostral middle frontal, medial/lateral orbitofrontal, frontal pole, and superior frontal lobe), and (5) a composite of regions thought to be early in accumulating Aβ (precuneus and posterior cingulate) (Palmqvist et al., 2017). 18F-florbetapir ROIs were expressed as SUVRs with a cerebellar reference region.

Methods to acquire and process tau (18F-flortaucipir) PET image data were described previously (Maass et al., 2017). Six tau ROI outcomes, corrected for partial-volume, were considered: (1) the medial temporal lobe (MTL) (amygdala, entorhinal and parahippocampal cortex), (2) the lateral temporal lobe (LTL) (inferior/middle/superior temporal lobe, banks of the superior temporal sulcus, transverse temporal lobe, temporal pole), (3) the medial parietal lobe (MPL) (isthmus cingulate, precuneus), (4) the lateral parietal lobe (LPL) (inferior/superior parietal lobe, supramarginal), (5) frontal lobe (pars, orbitofrontal and middle/superior frontal lobe), and (6) the occipital lobe (cuneus, lingual, pericalcarine, and lateral occipital lobe). 18F-flortaucipir ROIs were expressed as SUVRs with an inferior cerebellar grey matter reference region. Full details of PET acquisition and analysis can be found at http://adni.loni.usc.edu/methods/.

### Cognition

Cognitive measures assessed included the Mini-Mental State Examination (MMSE) as a measure of global cognition, and the Preclinical Alzheimer’s Cognitive Composite (PACC), as a measure of early AD-related cognitive changes. The PACC comprised the MMSE, the Logical Memory Delayed Word Recall from the Wechsler Memory Scale, the Alzheimer’s Disease Assessment Scale—Cognitive Subscale Delayed Word Recall, and the Trail Making Test part B (log transformed) (Donohue et al., 2017, 2014).

### Statistical Analysis

The aims of these analyses were to evaluate the relationship between the estimated TFAβ+ and CSF, PET, and cognitive responses. Because TFAβ+ was not directly observed, in a first step, linear mixed-effects models were fit to all available longitudinal global Aβ PET SUVR data to estimate subject-specific intercepts and slopes of Aβ pathology. Because Aβ slopes are unlikely to remain constant over long periods of time as subjects move toward and away from the Aβ threshold, natural splines (Hastie and Tibshirani, 1990) were used to estimate the nonlinear shape of the slopes with respect to baseline Aβ, using quantile regression. Rather than modeling the mean Aβ slope with respect to baseline Aβ, quantile regression provides a separate curve for each quantile, allowing the relationship between slope and intercept to differ depending on the location in the distribution of Aβ slope. For each subject, TFAβ+ was estimated by integrating over each subject’s quantile curve between the subject’s intercept and the threshold for Aβ-positivity (PET SUVR = 1.1). For example, for a subject with a baseline SUVR of 1.2 and a slope in the 0.6 quantile, TFAβ+ was taken to be the time it would take to go from SUVR = 1.1 to 1.2, using the slope estimates from the quantile curve. For incremental changes on the x-axis (baseline SUVR), the time required to travel the incremental distance is equal to distance/rate. Using the trapezoid rule (Atkinson, 1989), TFAβ+ is the sum of these incremental times spanning SUVR = 1.1 to 1.2. An example of calculating TFAβ+ is given in the top left panel of Figure 1.

**Figure 1.**
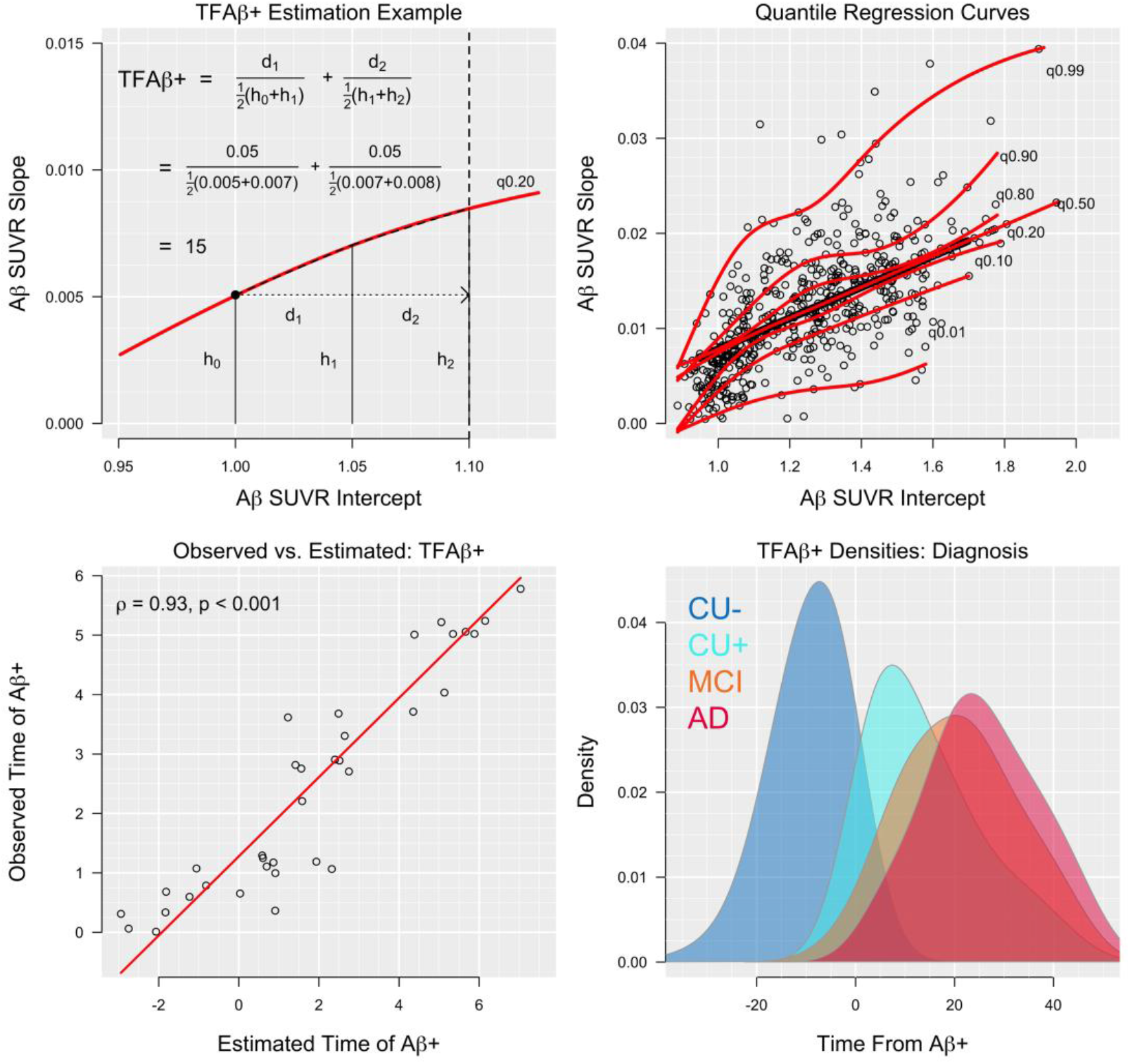
Observed vs. Estimated TFAβ+, Quantile Regression Curves and TFAβ+ Densities. Top left panel: an example of how TFAβ+ is estimated. Here we have a participant with an estimated intercept of SUVR = 1.00 and an accumulation rate (slope) of 0.005 SUVR/year. We want to calculate how long it will take for them to reach the 1.10 threshold. A slope of 0.005 SUVR/year puts this participant on the 0.20 quantile (20% percentile) curve. We know this participant must accumulate 0.10 SUVR to reach the threshold and we will assume they will continue to have an accumulation rate in the 0.20 quantile. Partitioning the curve into segments from SUVR = 1.00 to 1.10 and using the formula time = distance/rate, the time to cross each segment is calculated and summed. In the figure, only two segments are shown, but in the actual calculation, the curve is partitioned into a large number of segments. Assuming a linear rate increase within each segment (shown in the dashed black line along the red quantile curve), the time to travel the distance in the 1^st^ segment, from SUVR 1.00 to 1.05 is given by, time_1_ = d_1_ /rate_1_, where d1 is 0.05 and rate1 is the average rate in segment 1, which is ½(h_0_+h_1_), as shown in the panel. A similar calculation is done for segment 2 and the results are summed to give TFAβ+ = 15. Top right panel: quantile regression curves of Aβ PET slopes plotted against intercepts. Curves for several selected quantiles (0.01, 0.10, …, 0.99) are shown in red. Bottom left panel: observed time of Aβ+ plotted against estimated time of Aβ+. Bottom right panel: distributions of TFAβ+ for each group, Aβ-CU (CU-), Aβ+ CU (CU+), MCI, and AD are shown.

To evaluate the accuracy of the TFAβ+ estimates, we compared the observed times of Aβ+ to the estimated times of Aβ+ values in participants who were Aβ-at baseline and became Aβ+ during follow-up. Observed time of Aβ+ occurred in the interval between the last Aβ-scan and the first Aβ+ scan. The observed time was calculated as a weighted average of the two scan times, weighted proportionally toward the scan where the participant was closest to hitting the threshold. Observed and estimated values were compared in N=37 participants who crossed the threshold for Aβ+ and remained Aβ+ throughout follow-up.

Our analyses aim to model participants who are ostensibly on the AD trajectory and had calculable TFAβ+. Therefore, of the 963 participants with Aβ PET, we excluded N=16 participants with negative Aβ accumulation rates (negative rates were largely driven by one early high Aβ PET measure), we also excluded N=6 participants with low levels of Aβ and accumulation rates such that they were predicted to become Aβ+ later than 120 years of age (biomarker data from these subjects are included for visual comparisons in Figures 2–4, see Figure legends). We included subjects where the TFAβ+ metric indicated very early accumulation of Aβ, but for participants estimated to have become Aβ+ before age 40 (N=24, median estimated age at Aβ+ = 30, IQR: 24 to 34), we truncated TFAβ+ to age 40, based on previously described rates of Aβ-positivity in middle age (Jansen et al., 2015).

**Figure 2.**
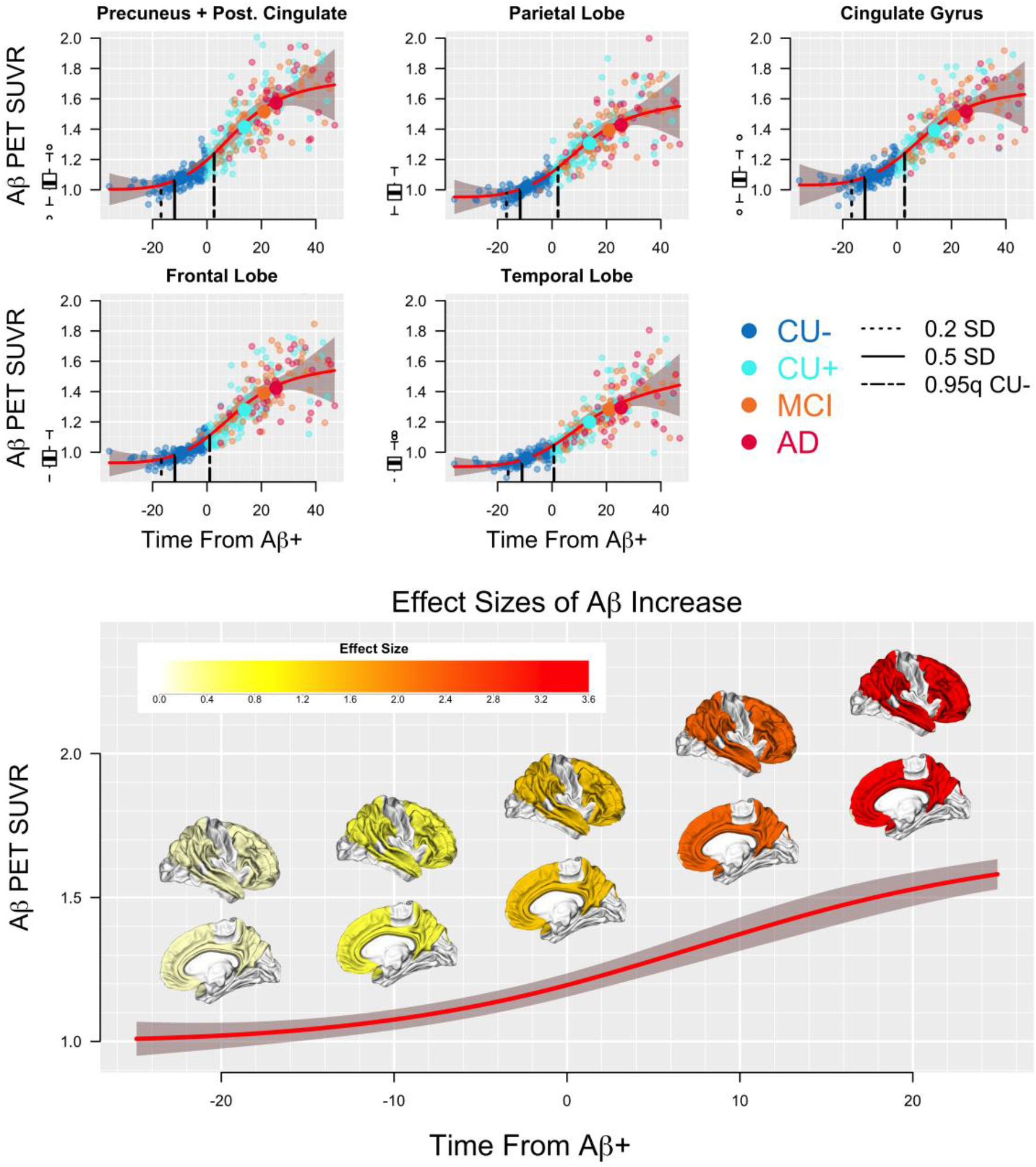
Regional Aβ PET. Aβ PET ROIs are plotted against TFAβ+. Effect sizes, depicting change points are shown as vertical dashed (0.2 SD, initial change) and solid (0.5 SD) lines. Regression curves (red) and corresponding 95% CIs (shaded grey) are shown. Mean values of the response are plotted against mean TFAβ+ for each of the four diagnosis groups (large symbols). The 0.95 quantile (approximately 1.65 SD if normally distributed) of the response for the CU-group is also shown (short/long dashed line). The 0.95 quantile (or 0.05 quantile for responses where low values are worse) of the biomarkers in CU-, provided for all responses to facilitate comparisons of when (in terms of TFAβ+) the average level of each response is no longer in the normal range. The boxplots to the left of each figure show the biomarker distribution in subjects that was determined to not be on the AD trajectory (including subjects where the model estimated them to become Aβ+ at over 120 years of age). Effect sizes of Aβ increase are shown in the bottom panel at TFAβ+ = −20, −10, 0, 10, and 20 years.

**Figure 3.**
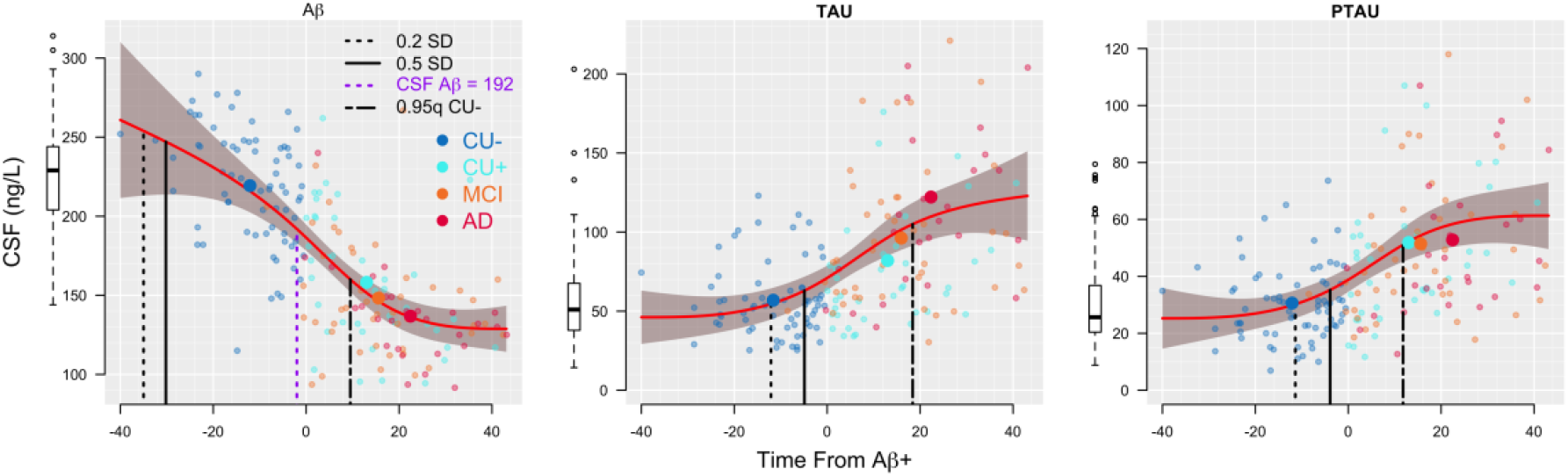
CSF Biomarkers. CSF biomarker responses are plotted against TFAβ+. Effect sizes, depicting change points are shown as vertical dashed (0.2 SD, initial change) and solid (0.5 SD) lines. Regression curves (red) and corresponding 95% CIs (shaded grey) are shown. Mean values of the response are plotted against mean TFAβ+ for each of the four diagnosis groups (large symbols). The 0.95 quantile (approximately 1.65 SD if normally distributed) of the response for the CU-group is also shown (short/long dashed line). The boxplots to the left of each figure show the biomarker distribution in subjects that were determined not to be on the AD trajectory (including subjects where the model estimated them to become Aβ+ over 120 years of age).

**Figure 4.**
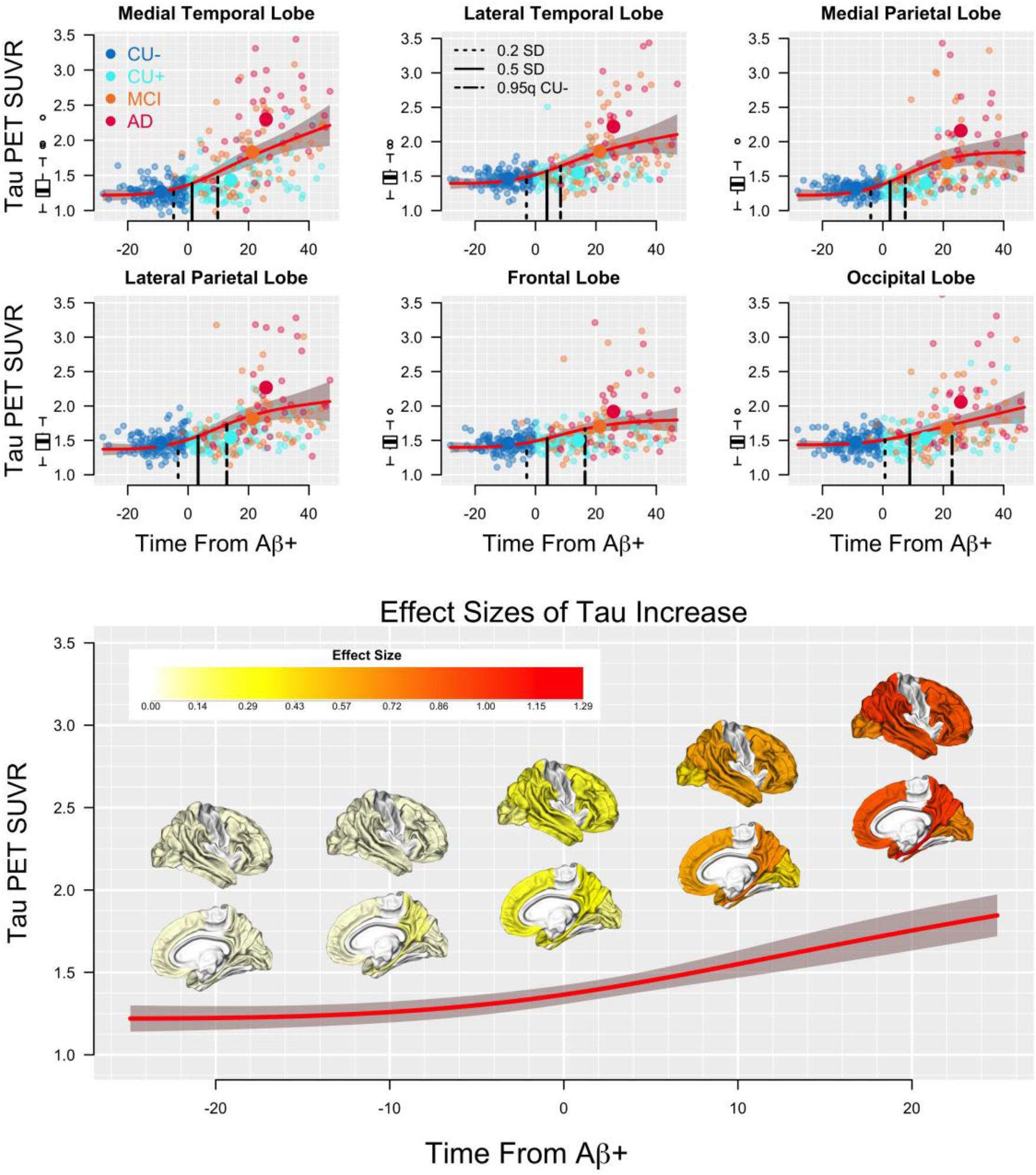
Regional Tau PET. Tau PET ROIs are plotted against TFAβ+. Effect sizes, depicting change points are shown as vertical dashed (0.2 SD, initial change) and solid (0.5 SD) lines. Regression curves (red) and corresponding 95% CIs (shaded grey) are shown. Mean values of the response are plotted against mean TFAβ+ for each of the four diagnosis groups (large symbols). The 0.95 quantile (approximately 1.65 SD if normally distributed) of the response for the CU-group is also shown (short/long dashed line). The boxplots to the left of each figure show the biomarker distribution in subjects that was determined to not be on the AD trajectory (including subjects where the model estimated them to become Aβ+ at over 120 years of age). Effect sizes of tau increase are shown in the bottom panel at TFAβ+ = −20, −10, 0, 10, and 20 years.

In the second step, the relationship between TFAβ+ and the responses was modeled using monotone penalized regression splines. Generalized cross-validation was used to tune the smoothing parameter (Wood, 1994). Cognitive responses were covaried for age, gender and education; CSF Aβ42, T-tau, P-tau and PET measures were covaried for age and gender.

In order to account for the uncertainty across steps 1 and 2, the entire process was repeated in 500 bootstrap samples to estimate 95% confidence intervals for the association between TFAβ+ and the responses.

Meaningful effect sizes of change of increase in pathology or decrease in cognition with respect to TFAβ+ were estimated. A Cohen’s d effect size of 0.2 SD was considered small, 0.5 SD was considered medium, and a 0.8 SD effect was considered large (Cohen, 1988). A 0.2 standard deviation (SD) change from the mean response at the longest times (least pathological) from Aβ-positivity was taken to be the initial point of meaningful change. A 0.5 SD change was also shown as a more substantial effect size of change. We also estimated change, 95% confidence intervals, and statistical significance of change for each response at TFAβ+ = 0, the time of Aβ-positivity, with bootstrap-estimated standard errors.

Baseline associations between demographics and TFAβ+ were assessed using Spearman correlation for age and education and the Wilcoxon rank-sum test for gender. Associations between diagnosis and demographics were assessed using Wilcoxon ranksum test for continuous variables and Fisher’s Exact test for categorical variables. All analyses were done in R v3.5.1 (www.r-project.org).

## Results

### Cohort Characteristics

Two-hundred and twenty-eight CU (128 Aβ- and 100 Aβ+), 70 Aβ+ MCI and 38 Aβ+ AD participants were included in the analysis. The diagnostic groups varied by mean age (CU-, 70.1 years old [SD=5.8]; CU+, 72.9 years old [SD=6.6]; MCI, 72.0 years old [SD=6.9]; AD, 74.5 years old [SD=7.2]; p<0.001). The diagnostic groups varied by sex (CU-, 60.2% female; CU+, 58.0% female; MCI, 41.4% female; AD, 47.4% female; p=0.05). The groups differed by mean years of education (CU-, 16.7 years [SD=2.4]; CU+, 16.9 years [SD=2.3]; MCI, 16.0 years [SD=2.5]; AD, 15.6 years [SD=2.5]; p=0.007). The groups varied by proportion of *APOE* ε4+ (CU-, 25.4%; CU+, 50.5%; MCI, 60.9%; AD, 54.3%; p<0.001).

### Aβ PET and Estimation of TFAβ+

TFAβ+ was estimated with a median of 3 (range: 1 to 5) Aβ PET scans per participant. The average time between first and last scan was 3.3 years (SD=2.9) and the average time between scans was 2.2 years (SD=0.8). Across diagnoses, TFAβ+ ranged from −35.9 to 47.0 years, where higher (positive) TFAβ+ values indicate more time spent with a significant Aβ burden. The average TFAβ+ was −9.3 years (SD=6.9) for CU-, 13.9 years (SD=11.2) for CU+, 21.1 years (11.7) for MCI, and 25.8 years (11.0) for AD participants (p<0.001, comparing CU+, MCI, and AD only). TFAβ+ was highly correlated with observed time of Aβ+ (ρ=0.93, p<0.001, bottom left panel of Figure 1). Higher TFAβ+ was significantly associated with older age (ρ=0.30, p<0.001), lower education (ρ=-0.15, p=0.01) and *APOE* ε4-positivity (mean TFAβ+ in *APOE* ε4-= 3.0 (SD=16.5) years and mean TFAβ+ in *APOE* ε4+ = 13.7 (SD=16.0) years, p<0.001). TFAβ+ was not associated with sex (mean TFAβ+ = 9.5 (SD=17.4) and 6.6 (SD=16.8) in males and females, respectively, p=0.13). Within-diagnosis TFAβ+ distributions are shown on the bottom right panel of Figure 1. Quantile curves of the relationship between Aβ intercepts and slopes are also shown in the top right panel of Figure 1, displaying the variation of acceleration of Aβ deposition over different levels of baseline Aβ.

### Regional Aβ PET

Five regional ROIs (precuneus + posterior cingulate, frontal lobe, cingulate gyrus, temporal and parietal lobes) are shown plotted against TFAβ+ in Figure 2. All 5 regions were estimated to reach a small, but meaningful (0.2 SD) increase in SUVR between 16-17 years before Aβ-positivity, i.e. TFAβ+ = 0. Effect sizes over the span of TFAβ+ are shown in Figure 2. At TFAβ+ = 0, all regions showed large, significant increases in SUVR (ΔSUVR ≥ 0.13, p≤0.01) with the precuneus + posterior cingulate composite showing the largest increase (ΔSUVR = 0.19, p < 0.01) and the temporal lobe showing the smallest (ΔSUVR = 0.13, p < 0.01). Effect sizes for all regions were large (>1) by the time of Aβ+. Table 1 summarizes the values of the responses at the longest times before Aβ+, i.e. the least pathological TFAβ+. Table 1 also shows the value and change of each response at the time of Aβ-positivity (TFAβ+ = 0), p-value and corresponding 95% confidence interval, the effect size of change of each response, and the 0.2 SD change point with respect to TFAβ+.

**Table 1.**
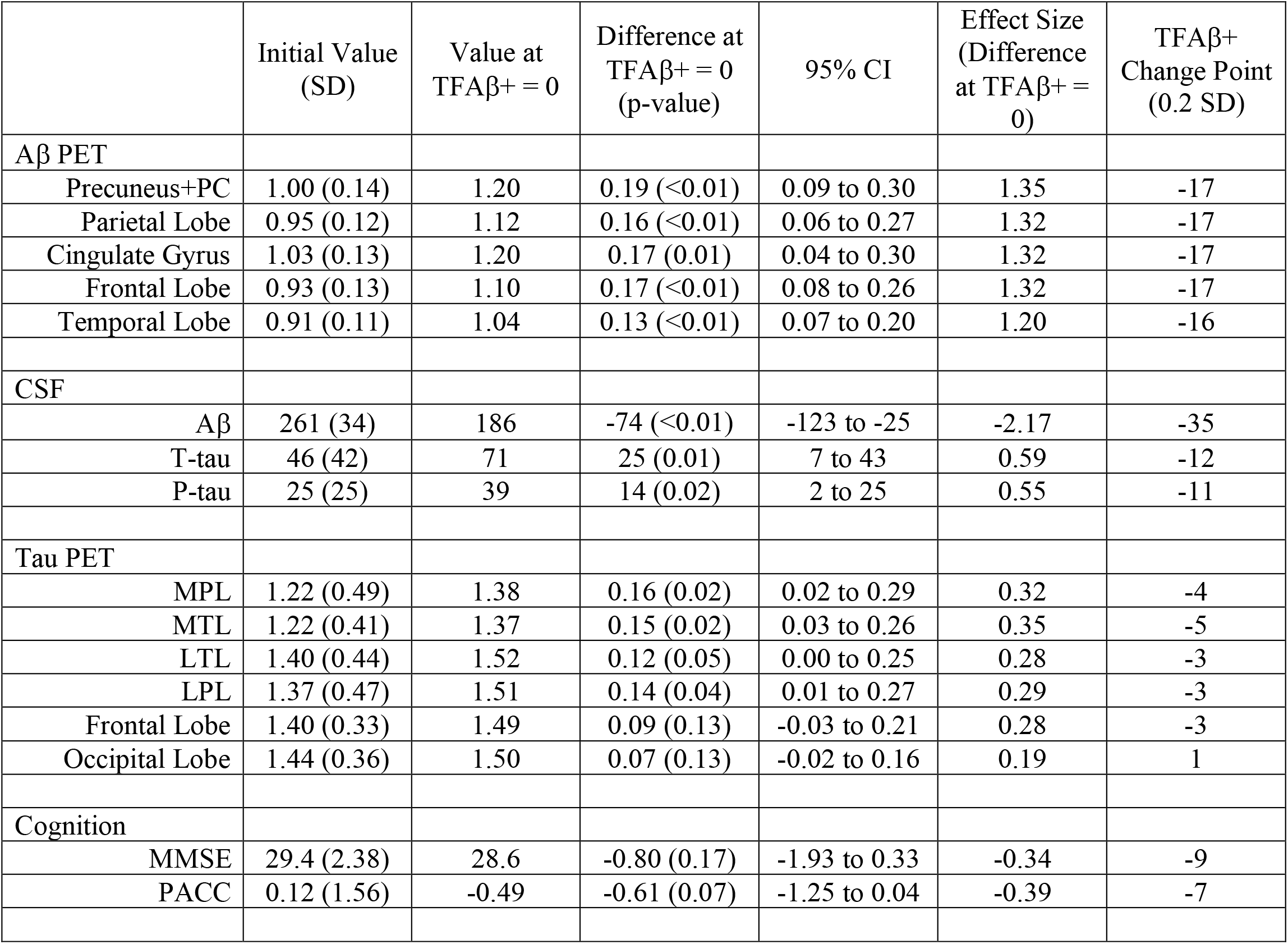
Initial values, effect sizes and change points

| | Initial Value<br>(SD) | Value at<br>TFAβ<sub>+</sub> = 0 | Difference at<br>TFAβ<sub>+</sub> = 0<br>(p-value) | 95% CI | Effect Size<br>(Difference<br>at TFAβ<sub>+</sub> =<br>0) | TFAβ<sub>+</sub><br>Change Point<br>(0.2 SD) |
| --- | --- | --- | --- | --- | --- | --- |
| **Aβ PET** |  |  |  |  |  |  |
| Precuneus+PC | 1.00 (0.14) | 1.20 | 0.19 (<0.01) | 0.09 to 0.30 | 1.35 | -17 |
| Parietal Lobe | 0.95 (0.12) | 1.12 | 0.16 (<0.01) | 0.06 to 0.27 | 1.32 | -17 |
| Cingulate Gyrus | 1.03 (0.13) | 1.20 | 0.17 (0.01) | 0.04 to 0.30 | 1.32 | -17 |
| Frontal Lobe | 0.93 (0.13) | 1.10 | 0.17 (<0.01) | 0.08 to 0.26 | 1.32 | -17 |
| Temporal Lobe | 0.91 (0.11) | 1.04 | 0.13 (<0.01) | 0.07 to 0.20 | 1.20 | -16 |
| **CSF** |  |  |  |  |  |  |
| Aβ | 261 (34) | 186 | -74 (<0.01) | -123 to -25 | -2.17 | -35 |
| T-tau | 46 (42) | 71 | 25 (0.01) | 7 to 43 | 0.59 | -12 |
| P-tau | 25 (25) | 39 | 14 (0.02) | 2 to 25 | 0.55 | -11 |
| **Tau PET** |  |  |  |  |  |  |
| MPL | 1.22 (0.49) | 1.38 | 0.16 (0.02) | 0.02 to 0.29 | 0.32 | -4 |
| MTL | 1.22 (0.41) | 1.37 | 0.15 (0.02) | 0.03 to 0.26 | 0.35 | -5 |
| LTL | 1.40 (0.44) | 1.52 | 0.12 (0.05) | 0.00 to 0.25 | 0.28 | -3 |
| LPL | 1.37 (0.47) | 1.51 | 0.14 (0.04) | 0.01 to 0.27 | 0.29 | -3 |
| Frontal Lobe | 1.40 (0.33) | 1.49 | 0.09 (0.13) | -0.03 to 0.21 | 0.28 | -3 |
| Occipital Lobe | 1.44 (0.36) | 1.50 | 0.07 (0.13) | -0.02 to 0.16 | 0.19 | 1 |
| **Cognition** |  |  |  |  |  |  |
| MMSE | 29.4 (2.38) | 28.6 | -0.80 (0.17) | -1.93 to 0.33 | -0.34 | -9 |
| PACC | 0.12 (1.56) | -0.49 | -0.61 (0.07) | -1.25 to 0.04 | -0.39 | -7 |

### CSF

CSF responses are plotted against TFAβ+ in Figure 3. A 0.2 SD drop in CSF Aβ42 was estimated to occur 35 years before Aβ-positivity (TFAβ+ = −35). At TFAβ+ = 0, CSF Aβ42 showed a very large effect size (ΔAβ42 = −74 ng/L, p<0.01, effect size = - 2.17). At TFAβ+ = −2, or two years before Aβ-positivity, the population curve passes through a previously published CSF Aβ42 threshold for Aβ-positivity (192 ng/L) (Shaw et al., 2009).

A 0.2 SD increase in CSF T-tau and P-tau was estimated to occur 11-12 years before the time of Aβ-positivity (TFAβ+ = −12 and −11, respectively). At TFAβ+ = 0, significant increases of medium effect size of T-tau (ΔT-tau = 25 ng/L, p=0.01, effect size = 0.59) and P-tau (ΔP-tau = 14 ng/L, p=0.02, effect size = 0.55) were observed.

### Tau PET

Six regional ROIs (MTL, LTL, MPL, LPL, frontal and occipital lobes) are shown plotted against TFAβ+ in Figure 4. Five of the six regions were estimated to reach a 0.2 SD increase in SUVR 3-5 years before Aβ-positivity, with the occipital lobe reaching a 0.2 SD increase one year after Aβ-positivity. Effect sizes over the span of TFAβ+ are shown in Figure 4. At TFAβ+ = 0, four regions (MTL, LTL, MPL, LPL) showed significant small increases in SUVR (ΔSUVR ≥ 0.12, p≤0.05) with the MTL showing the largest effect size (0.35). The frontal and occipital lobes did not increase significantly by TFAβ+ = 0 (ΔSUVR = 0.09, 0.07, respectively, p=0.13). Estimates are summarized in Table 1.

### Cognition

Cognitive measures are shown in Figure 5. The MMSE showed a 0.2 SD drop nine years before Aβ-positivity, followed by the PACC seven years before Aβ-positivity. Neither measure decreased significantly by the time of Aβ-positivity, although the PACC was borderline (ΔMMSE = −0.80, p=0.17, effect size=-0.34; ΔPACC = −0.61, p=0.07, effect size=-0.39). Summary curves and 0.2 SD change points are shown in Figure 6.

**Figure 5.**
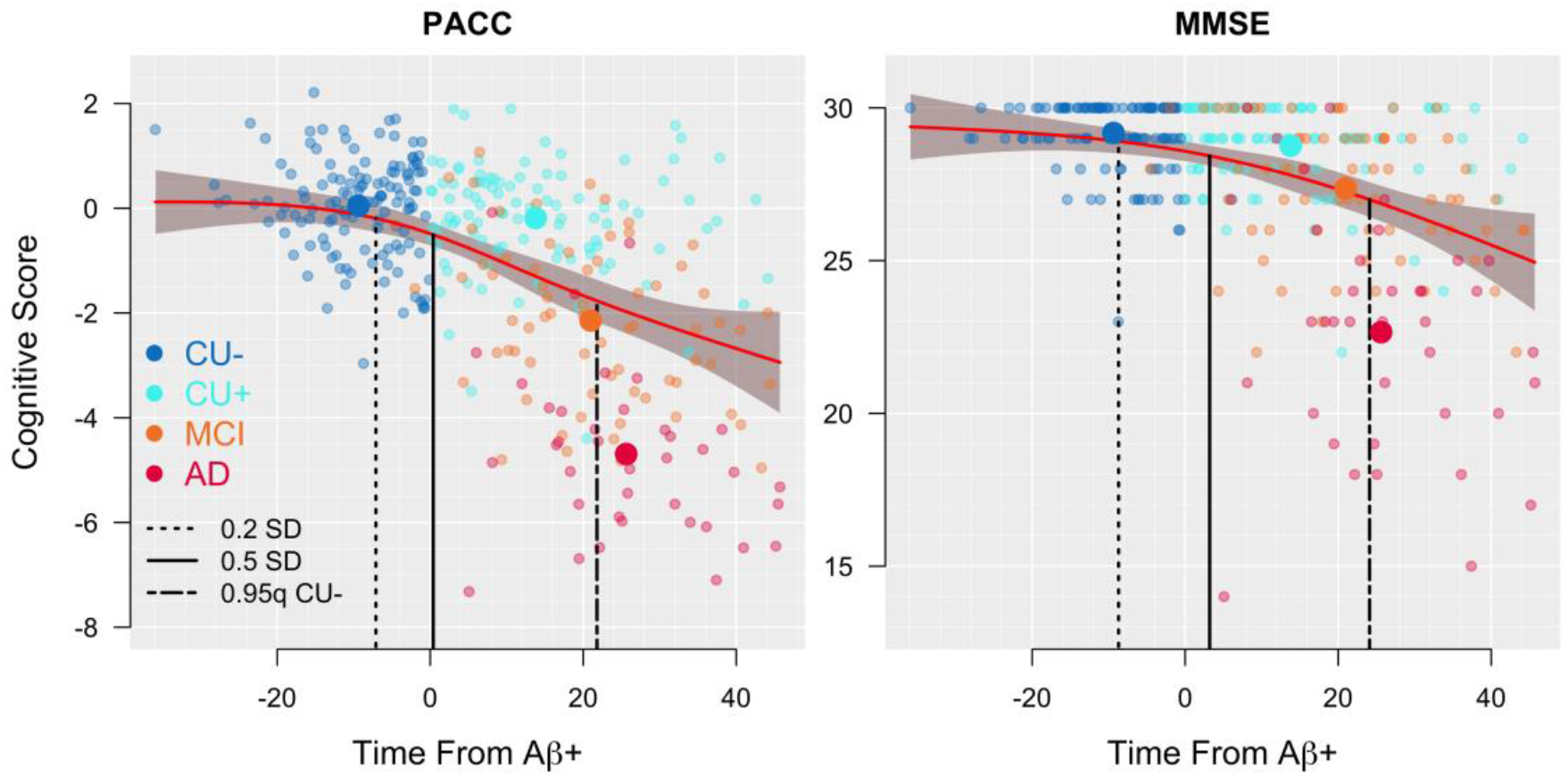
Cognition. MMSE and PACC scores are plotted against TFAβ+. Effect sizes, depicting change points are shown as vertical dashed (0.2 SD, initial change) and solid (0.5 SD) lines. Regression curves (red) and corresponding 95% CIs (shaded grey) are shown. Mean values of the response are plotted against mean TFAβ+ for each of the four diagnosis groups (large symbols). The 0.05 quantile (approximately −1.65 SD if normally distributed) of the response for the CU-group is also shown (short/long dashed line).

**Figure 6.**
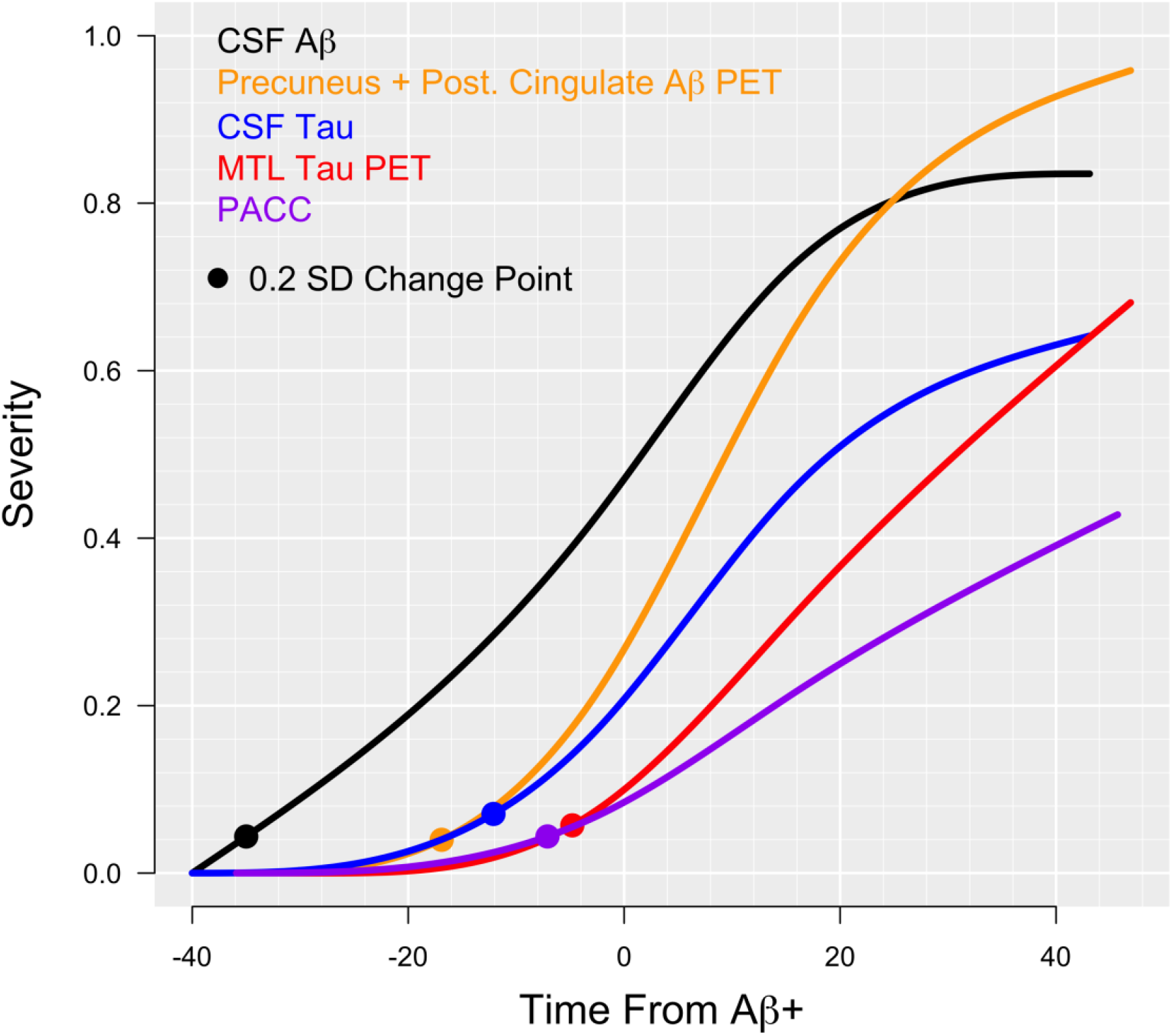
Summary Curves. Summary curves are shown for all modalities on a scale from zero to one. Responses are scaled such that zero is the least pathological point for each response and one is the mean response in the AD participants. The initial effect, defined by 0.2 SD change points are plotted.

## Discussion

Several biological processes develop over time in sporadic AD, including accumulation of Aβ and tau across wide areas of the brain, as well as cognitive decline. Based on the amyloid cascade hypothesis, a relevant overarching time scale of the disease processes could be based on the development of Aβ pathology (Koscik et al., 2020). We have therefore integrated Aβ PET level and rate of change information to place each individual on a pathological timeline. This timeline can then be used to estimate the time of downstream events in the amyloid cascade. We estimated several major milestone events of AD progression including a small drop in CSF Aβ42 35 years before Aβ-positivity and a small increase in regional Aβ PET deposition 17 years before Aβ-positivity. Using the biomarkers tested here, the first changes in CSF Aβ42 may define the onset of AD. Small increases in tau pathology were estimated to occur 11-12 years before Aβ-positivity, as measured by CSF and 5 years before, as measured by PET. More substantial and statistically significant increases in CSF as well as temporoparietal tau PET were detected by the time of Aβ-positivity. Small effects of cognitive dysfunction occurred 7-9 years before Aβ-positivity, coinciding with previous reports (Insel et al., 2017). These findings provide a general time scale for initial changes in sporadic AD, which may inform clinical trials aimed at specific stages of the disease.

A 0.2 SD difference, a small, but meaningful increase in levels of CSF tau and temporoparietal lobe tau are observed years before the current threshold for Aβ-positivity. In the context of secondary prevention trials where Aβ-positivity at current thresholds is required for study inclusion, tau levels in these participants would already have been increasing for several years, likely more. The finding that temporoparietal tau starts to increase prior to other regions is in accordance with 18F-flortaucipir studies on other populations. Cross-sectional studies showed early tau deposition in cognitively healthy elderly (with or without significant Aβ pathology) in temporal and medial parietal regions, most dominant in entorhinal and parahippocampal cortex, the amygdala and inferior temporal cortex. Longitudinal studies further suggest that cognitively healthy elderly accumulate tau in the medial temporal and medial parietal lobe, while (Aβ positive) AD dementia patients increased in tau primarily in the frontal lobe (Harrison et al., 2018). The spread of tau beyond the MTL to the parietal lobe and other regions may be a critical milestone in the progression of AD. The early changes observed in the MPL in this study coincide with a recent report of the earliest tau deposition found in medial parietal regions (precuneus and isthmus cingulate) in autosomal dominant AD (Gordon et al., 2019). Considering that a 0.2 SD increase in MPL tau can potentially be detected several years before Aβ-positivity (Figure 4), these data support the use of primary prevention trials against Aβ where treatment is initiated years before the current threshold for Aβ-positivity, if treatment efficacy relies on early intervention, prior to the development of tau pathology.

The initial descent in cognitive performance is estimated to occur 7-9 years before becoming Aβ+ (Figure 5). Reduced cognitive performance has repeatedly been shown to be associated with elevated levels of Aβ (Baker et al., 2017; Donohue et al., 2017; Insel et al., 2017, 2016), even within the subthreshold range (Landau et al., 2018), in cognitively unimpaired individuals. The result that CSF tau measures started to change between regional Aβ and cognition in this study is in accordance with the theory that cognitive impairment in AD is caused primarily by tau pathology. This is also in line with other recent studies which show that cognitive impairment is more strongly related to accumulation of tau than to Aβ (Ossenkoppele et al., 2019), and that both tau and Aβ appear necessary for cognitive decline (Sperling et al., 2019). The ordering of the responses coincides with the magnitude of the effect sizes at the time of Aβ-positivity (Table 1), suggesting that initial changes in the responses continue to change in parallel through to the time of Aβ-positivity, without any major differences in acceleration.

In their 2018 draft guidance, the FDA indicated that because it is highly desirable to intervene as early as possible in AD, it follows that patients with characteristic pathophysiologic changes of AD but no subjective complaint, functional impairment, or detectable abnormalities on sensitive neuropsychological measures are an important target for clinical trials (Food and Drug Administration, 2018). If the spread of tau to the lateral temporal and parietal lobes becomes a defining characteristic of pathophysiological change in AD, the window to intervene as early as possible may shift to years before the current threshold for Aβ-positivity. It is possible that early accelerations of tau may have contributed to recent failures of anti-Aβ treatments in phase III clinical trials on Aβ-positive patients (Egan et al., 2018; Honig et al., 2018). Although selecting subjects that are Aβ-positive ensures that only AD patients are included in trials, the use of conservative thresholds to define Aβ-positivity may bias trial populations toward individuals where tau pathology has already accumulated, causing downstream injuries independent of Aβ, reducing the efficacy of anti-Aβ treatments.

This study has several limitations. Tau PET data were available for only a subsample of the data, limiting comparisons to a small cross-section of the full ADNI data set. More data, especially longitudinal data in participants in the earliest stages of Aβ changes, will be required for more precise change point estimates. These analyses lack the power and precision to place the temporal and parietal tau regions in a particular order with confidence, but instead demonstrate that temporoparietal tau increases years before Aβ-positivity. The ADNI CU, MCI and AD cohorts are also age matched. The AD patients, on average, have dementia by age 75, while the participants in the CU cohort who may eventually develop AD, are unlikely to do so for many years, possibly decades. By design, these cohorts with age matched groups are therefore on systematically different disease trajectories with respect to age. If earlier onset is associated with a more aggressive form of the disease, then the AD cohort may have the most aggressive form while the CU cohort, the least aggressive. If the developing Aβ pathology in the ADNI CU-cohort represents a less aggressive disease process compared with a more typical AD process, the estimates reported here could be conservative and biased toward later time estimates for downstream events. The ADNI MCI cohort may represent a more typical trajectory with respect to downstream events along the Aβ pathological timeline. These differences in disease trajectories are apparent from the cohort estimates in Figures 2–5. Additionally, the change point estimates are influenced by both biological variation and measurement error, which varies from marker to marker. Change points in measures with high variability in the “normal” range and excess measurement error may require additional biological change to detect, despite an earlier, real increase in pathology. ADNI participants are highly educated on average, reducing generalizability to some degree. The associations between increasing Aβ pathology and downstream changes, including increased tau pathology reported here do not imply causality. It remains unknown whether and to what degree downstream pathological changes can be directly attributed to the accumulation of Aβ. Only studies with experimental interventions against Aβ-pathology, with clear verification of target engagement, can be used to show causal relationships between Aβ-deposition and putative downstream events.

Longitudinal information is required to evaluate how quickly an individual’s pathophysiological changes are occurring and to accurately characterize their disease trajectory. Analyses limited to a cross-sectional evaluation of Aβ status are naïve to the time spent with a significant Aβ burden. Incorporating longitudinal information facilitates the estimation of the time-course of downstream events such as the spread of tau and the onset of subtle cognitive dysfunction. As the technology to measure AD pathology becomes more cost effective and noninvasive, such as plasma measures of Aβ or tau (Janelidze et al., 2020; Mielke et al., 2018; Palmqvist et al., 2019; Schindler et al., 2019), longitudinal evaluations in the context of trial-ready cohorts may greatly improve early diagnosis and expedite the execution of clinical trials in early AD.

## Acknowledgements

Data collection and sharing for this project was funded by the Alzheimer’s Disease Neuroimaging Initiative (ADNI) (National Institutes of Health Grant U01 AG024904). ADNI is funded by the National Institute on Aging, the National Institute of Biomedical Imaging and Bioengineering, and through generous contributions from the following: Alzheimer’s Association; Alzheimer’s Drug Discovery Foundation; BioClinica, Inc.; Biogen Idec Inc.; Bristol-Myers Squibb Company; Eisai Inc.; Elan Pharmaceuticals, Inc.; Eli Lilly and Company; F. Hoffmann-La Roche Ltd and its affiliated company Genentech, Inc.; GE Healthcare; Innogenetics, N.V.; IXICO Ltd.; Janssen Alzheimer Immunotherapy Research & Development, LLC.; Johnson & Johnson Pharmaceutical Research & Development LLC.; Medpace, Inc.; Merck & Co., Inc.; Meso Scale Diagnostics, LLC.; NeuroRx Research; Novartis Pharmaceuticals Corporation; Pfizer Inc.; Piramal Imaging; Servier; Synarc Inc.; and Takeda Pharmaceutical Company. The Canadian Institutes of Health Research is providing funds to support ADNI clinical sites in Canada. Private sector contributions are facilitated by the Foundation for the National Institutes of Health (www.fnih.org). The grantee organization is the Northern California Institute for Research and Education, and the study is coordinated by the Alzheimer’s Therapeutic Research Institute at the University of Southern California, San Diego. ADNI data are disseminated by the Laboratory for Neuro Imaging at the University of Southern California. Data used in preparation of this article were obtained from the Alzheimer’s Disease Neuroimaging Initiative (ADNI) database (adni.loni.usc.edu). As such, the investigators within the ADNI contributed to the design and implementation of ADNI and/or provided data but did not participate in analysis or writing of this report. A complete listing of ADNI investigators can be found at: http://adni.loni.usc.edu/wp-content/uploads/how_to_apply/ADNI_Acknowledgement_List.pdf.

This research was also supported by The Wallenberg Center for Molecular Medicine at Lund University, the Knut and Alice Wallenberg foundation, The Medical Faculty at Lund University, Region Skåne, the Skåne University Hospital Foundation, the Swedish Research Council, the Swedish Alzheimer Foundation, the Swedish Brain Foundation, the Swedish Medical Association, the Konung Gustaf V:s och Drottning Victorias Frimurarestiftelse, the Greta and Johan Kock Foundation, the Thelma Zoega Foundation, the Gyllenstiernska Krapperupsstiftelsen, the Magnus Bergwall Foundation, the Bundy Academy, the Marianne and Marcus Wallenberg foundation, and the Strategic Research Area MultiPark (Multidisciplinary Research in Parkinson’s disease) at Lund University. The funding sources had no role in the design and conduct of the study, in the collection, analysis, interpretation of the data or in the preparation, review or approval of the manuscript.

## Competing Interests

Mr. Insel, Dr. Berron and Dr. Donohue report no competing interests.

Dr. Mattsson-Carlgren has been a consultant for ADNI.

Dr. Hansson has acquired research support (for the institution) from Roche, GE

Healthcare, Biogen, AVID Radiopharmaceuticals and Euroimmun. In the past 2 years, he has received consultancy/speaker fees (paid to the institution) from Biogen and Roche.

